# Regulating Soil Bacterial Diversity, Enzyme Activities and Community Composition Using Residues from Golden Apple Snails

**DOI:** 10.1101/692822

**Authors:** Jiaxin Wang, Xuening Lu, Jiaen Zhang, Guangchang Wei, Yue Xiong

## Abstract

Golden apple snails (GAS) have become a serious pest for agricultural production in Asia. A sustainable method for managing GAS is urgently needed, including potentially using them to produce commercial products. In this study, we evaluate the effects of GAS residues (shell and meat) on soil pH, bacterial diversity, enzyme activities, and other soil characteristics. Results showed that the amendment of GAS residues significantly elevated soil pH (to near-neutral), total organic carbon (TOC) (by 10-134%), NO_3_-N (by 46-912%), NH_4_-N (by 18-168%) and total nitrogen (TN) (by 12-132%). Bacterial diversity increased 13% at low levels of amendment and decreased 5% at high levels, because low-levels of GAS residues increased soil pH to near-neutral, while high-levels of amendment substantially increased soil nutrients and subsequently suppressed bacterial diversity. The dominant phyla of bacteria were: Proteobacteria (about 22%), Firmicutes (15-35%), Chloroflexi (12%-22%), Actinobacteria (8%-20%) Acidobacteria, Gemmatimonadetes, Cyanobacteria and Bacterioidetes. The amendment of GAS residues significantly increased the relative abundance of Firmicutes, Gemmatimonadetes, Bacterioidetes and Deinococcus-Thermus, but significantly decreased the relative abundance of Chloroflexi, Actinobacteria, Acidobacteria, Cyanobacteria and Planctomycetes. Our results suggest that GAS residues treatment induces a near-neutral and nutrient-rich soil. In this soil, soil pH may not be the best predictor of bacterial community composition or diversity; rather soil nutrients (*ie*., NH_4_-N and NO_3_-N) and soil TOC showed stronger correlations with bacterial community composition. Overall, GAS residues could replace lime for remediation of acidic and degraded soils, not only to remediate physical soil properties, but also microbial communities.

**Importance:** The wide spreading golden apple snail (GAS) is a harmful pest to crop productions and could result in soil and air pollutions after death. In the previous study, we developed a biocontrol method: adding GAS residues to acidic soil to mitigate the living GAS invasion and spread, improve soil quality, and reduce soil and air pollution. However, the effects of GAS residues amendment on bacterial diversity and community still remain unclear. This study provided insights into bacterial diversity and community compositions to facilitate the evaluation of GAS residues application.

## 1 Introduction

Invasive golden apple snails (GAS) *Pomacea canaliculata (Lamarck*) have become a serious pest for agricultural production in Asia (1). A series of control methods have been developed (2, 3); the most widely used is chemical control by molluscicides (4), which, could harm environmental and human health (5). A better method for sustainable management is urgently needed, potentially by using them to produce commercial products.

Studies have reported that GAS contains abundant CaCO_3_ and proteins, and could be used as feed for livestock such as pigs and ducks (6). However, only small GAS can be eaten by ducks or pigs, because the hard shells of large adult GAS make them unpalatable. The CaCO_3_ in GAS can also neutralize acidic soils, similar to lime. However, lime tends to lead to soil compaction, Si and P deficiency, and reduced soil microbial biomass and diversity.

Anthropogenic N inputs to terrestrial ecosystems have increased three- to five-fold over the past century (7). High levels of N fertilization can drive soil acidification both directly and indirectly (8). J. H. Guo et al. (8) found that anthropogenic acidification driven by N fertilization is at least 10 to 100 times greater than that associated with acid rain. The application and deposition of N is expected to continue to increase (9).

To alleviate soil acidification and control the invasion of the GAS, we have proposed using powered residues from GAS to mitigate soil acidification (unpublished data). The application of GAS residues can significantly increase soil pH and nitrate nitrogen (NO_3_-N) both at a high and low amendment levels, and can increase soil total organic carbon (TOC), total nitrogen (TN), ammonium nitrogen (NH_4_-N) and NO_3_-N compared with controls and with liming, which has no significant effect on soil nutrients (unpublished data). Although our previous research has indicated that the addition of GAS residues could significantly increase soil microbial biomass and regulate microbial community structure, the mechanisms by which GAS residues regulate microbial community composition, relative abundance and diversity remain unknown.

Previous studies have proposed soil pH and N input as the main predictors of soil microbial diversity in soils (10-12). However, the responses of soil microbes to elevated N inputs and pH are inconsistent. Numerously studies have revealed that N addition led to significant reductions in soil microbial activity (13), diversity (14) and community composition (15) because of increases in C sequestration and/or decreased soil respiration rates (13). Some studies have suggested that neutral soils support greater bacterial diversity than do acidic soils (10, 11). However, some researchers have suggested the opposite—that forest soils with lower pH support greater microbial diversity than agricultural soils with higher pH values (16).

Soil extracellular enzymes play key roles in biological soil processes, specifically in the degradation of soil organic compounds and the mineralization and recycling of nutrients related to C, N, P and S (17). Environmental factors, including water (18), salinity (18), pollution (19), soil nutrients (20), temperature (21) and soil pH can affect the activities of extracellular enzymes directly or indirectly.

Here, we conducted a series of greenhouse experiments amending GAS residues and lime to acidic and degraded soils. We hypothesize that GAS residues and lime—both of which increase soil pH—may differ in their regulation of microbial community structure, diversity and microbial enzyme activities. The objectives of this study were: (1) to explore bacterial community composition and diversity in soils neutralized with powered GAS residues, (2) to determine key factors controlling the composition of bacterial communities, (3) and assess differences in bacterial communities in soils treated with GAS residues and soils treated with lime.

## 2 Materials and methods

### 2.1 Testing materials

Golden apple snails (GAS) were collected from the paddy fields at the Xin Tang (Ning Xi) Research and Educational Station of South China Agricultural University, which is located in Zengcheng City, Guangdong Province, China. The snails were washed and frozen at −40°C in a freezer for 24h. Then the dead snails were dried, grounded into powder and stored in a desiccator. A slightly acidic soil was also collected from the paddy fields with the pH levels ranging from 6.25 to 6.53. The soil predominately consists of medium (38%) and fine (22%) sand, silt (36%) and clay (4%) and has 16.38 g kg^-1^ of TOC, 2.20 g kg^-1^ of TN and 0.58 g kg^-1^ of total phosphorus (TP).

### 2.2 Experimental design

Experiments were conducted in a greenhouse on the campus of South China Agricultural University. We implemented three treatments: (1) the control treatment with soil only (CK); (2) soil amended with GAS residues (*i.e*., shell and meat) (SR); and (3) soil amended with lime (SL). Each treatment had six levels of amendment: 0.5, 1, 2.5, 25, 50, and 100 g kg^-1^. Amendments were homogenized with soil in each treatment and carefully packaged in a polyvinyl chloride (PVC) column with a diameter of 180 mm and a height of 260 mm. A filter paper was mounted at the bottom of the column to prevent loss of soil or GAS residues. A base plate was also placed at the bottom of the column to position the column. About 400 ml of deionized water (pH = 7.0±0.1) were sprayed to the column each week to prevent the column from drying. Each treatment was triplicated in this study. After 120 days incubation, the soil samples were collected and stored at 4 °C for about 4h.

### 2.3 Soil analysis

Soil samples were homogenized by hand and passed through a 2mm sieve to remove rocks, roots and organic residues, and then divided evenly into three subsamples. These three subsamples were treated as follows: the first subsamples were stored at room temperature (about 22 °C) for about 4h, the second subsamples were stored at 4 °C, and the third subsamples were stored at −20 °C for further analysis. The room temperature (first) subsamples were air dried, grounded and analyzed for soil pH, TOC, TN, available phosphorous (AP), ammonium nitrogen (NH_4_–N) and nitrate nitrogen (NO_3_–N). The second subsamples were analyzed for soil gravimetric moisture and extracellular enzyme activity. The third subsamples were freeze-dried and grounded into power and passed through a 0.25 mm sieve for high-throughput sequencing analysis.

Soil pH was measured from fresh soil slurries (1 g of soil: 2.5 ml of deionized water) using a handheld multiparameter meter (SX-620, SAN XIN, China). Approximately 20 mg of each powered sample was analyzed for TOC and TN using a C analyzer (TOC-VCSH, Shimadzu Corp., Kyoto, Japan). Concentrations of NH_4_–N were analyzed using a UV-vis spectrophotometer at a wavelength of 420 nm (22). Concentrations of NO_3_–N were determined using a UV-vis spectrophotometer applying double wavelength of 275 nm and 220 nm (22). Concentrations of AP were analyzed using the molybdenum-antimony anti-spectrophotometric method (23).

### 2.4 Extracellular enzyme assay

Soil enzyme activities, including β–1,4–glucosidase (BG), acid phosphatase (ACP), β–1,4–N–acetylglucosaminidase (NAG) and β–D–cellobiosidase (cellulose degradation; CB), were measured using fluorometry as described in C. W. Au - Bell et al. (24), with minor modifications. In brief, 1 g of soil was weighted and dissolved in a porcelain evaporating dish with 125 ml of acetate-acetate buffer solution and blended for 30 min on a magnetic stirrer (JB-3, Ronghua, Jianfsu) at 200 rpm. Soil buffer solution (200 μl) was sucked from the dish and injected into a deep-well plate (Labtide, Greystone Biosciences LLC, USA), and then 50 μl each of buffer, substrate and 4–Methylumbelliferyl (MUB) was pipetted and injected into deep-well plates, representing a blank well, sample well and quench well, respectively. We used 200 μl of buffer plus 50 μl of substrate as the negative well and 200 μl buffer plus 50 μl of MUB as the reference standard well. Incubation was conducted in a constant temperature incubator (RXZ, Dongqi, Ningbo, China) at 37 °C for 3 h. After incubation, the sample was centrifuged (Eppendorf, USA) at 2900 rpm for 3 min, and the supernate was pipetted and injected into a black flat-bottomed 96-well microplate for fluorescence determination in a microplate reader (SYNERGY H1, BioTek, USA) at the excitation wavelength of 365 nm and emission wavelength of 450 nm.

### 2.5 Bacterial community and diversity analysis

Bacterial community structure and diversity in soils were determined by 16S rRNA. Soil DNA was extracted with the E.Z.N.A.^®^ soil DNA Kit (Omega Bio-tek, Norcross, GA, USA.) in accordance with manufacturer’s instructions. The extracted DNA was quantified by spectrophotometry (Nanodrop 2000, Thermo Scientific, UAS) and stored at −20 °C. Polymerase chain reaction (PCR) was carried out using the universal primer set 338F (5’-ACTCCTACGGGAGGCAGCAG-3’) and 806R (5’-GGACTACHVGGGTWTCTAAT-3’) by thermocycler PCR system (GeneAmp 9700, ABI, USA). PCR was performed to amplify 1 μl of template DNA in a 20-μl reaction system containing 4 μl of 5 × FastPfu Buffer, 2 μl of 2.5 mM dNTPs, 0.8 μl of each primer (5 μM), 0.4 μl of FastPfu Polymerase and 10 ng of template DNA. Amplification was performed in triplicate as follows: 95 °C for 3 min; 27 cycles of 95 °C for 30 s, 30s for annealing at 55 °C, and 45s for elongation at 72 °C; and a final extension at 72 °C for 10 min. PCR reactions were performed in triplicate 20 μl mixture containing 4 μl of 5 × FastPfu Buffer, 2 μl of 2.5 mM dNTPs, 0.8 μl of each primer (5 μM), 0.4 μl of FastPfu Polymerase and 10 ng of template DNA. The resulting PCR products were extracted from a 2% agarose gel and further purified using the AxyPrep DNA Gel Extraction Kit (Axygen Biosciences, Union City, CA, USA) and quantified using QuantiFluor TM -ST (Promega, USA) according to the manufacturer’s protocol. Purified amplicons were pooled in equimolar and paired-end sequenced (2 × 300) on an Illumina MiSeq platform (Illumina, San Diego, USA) according to the standard protocols by Majorbio Bio-Pharm Technology Co. Ltd. (Shanghai, China). Raw fastq files were demultiplexed, quality-filtered by Trimmomatic and merged by FLASH with the following criteria: (i) The reads were truncated at any site receiving an average quality score < 20 over a 50 bp sliding window. (ii) Primers were exactly matched allowing 2 nucleotide mismatching, and reads containing ambiguous bases were removed. (iii) Sequences with overlaps longer than 10 bp were merged according to their overlap sequences. Operational taxonomic units (OTUs) were clustered with 97% similarity cutoff using UPARSE (version 7.1 http://drive5.com/uparse/) and chimeric sequences were identified and removed using UCHIME. The taxonomy of each 16S rRNA gene sequence was analyzed by RDP Classifier algorithm (http://rdp.cme.msu.edu/) against the Silva (SSU123) 16S rRNA database using confidence threshold of 70%.

### 2.6 Data analysis

Analysis of Variance (ANOVA) was performed using SPSS. Differences in microbial communities were tested using ANOSIM with 9,999 permutations. A non-metric multidimensional scaling (NMDS) ordination to illustrate the clustering of bacterial community composition variation was conducted using the Vegan software based on the Bray-Curtis distance of genus. We used spearman correlations to identify significant relationships between soil parameters (involved in six amendment levels of GAS residues and five amendment levels of lime treatments) and the most abundant phyla and genera. To evaluate bacterial diversity in amended soils, Shannon index (25) was applied. To compare microbial communities and identify specialized communities in samples, we used the LEfSe tool (26). Statistical analysis was performed only from the phylum to the genus level to simplify the computation. Non-metric multidimensional scaling (NMDS) of soil bacteria community composition based on Bray-Curtis distances linear regression after amending with lime and GAS residues were conducted to explore the main factors affected soil bacterial community compositions.

## 3 Results and discussion

### 3.1 Amendment effects on soil properties

Amendments of GAS residues and lime resulted in increased soil pH (Fig. 1a). The addition of 1.0-2.5 g kg^-1^ GAS residue increased soil pH to neutral (7.0) and additional amendments further increasing pH. Even the smallest amendment of lime (0.5 g kg^-1^) changed soil pH sharply from acidic to light alkaline (pH > 7.8) (Fig. 1a). The effect of GAS residues on pH may have been mitigated by the decomposition of proteins (from the snail meat) into amino acids and glucose by soil microbes (27).

**Fig. 1.**
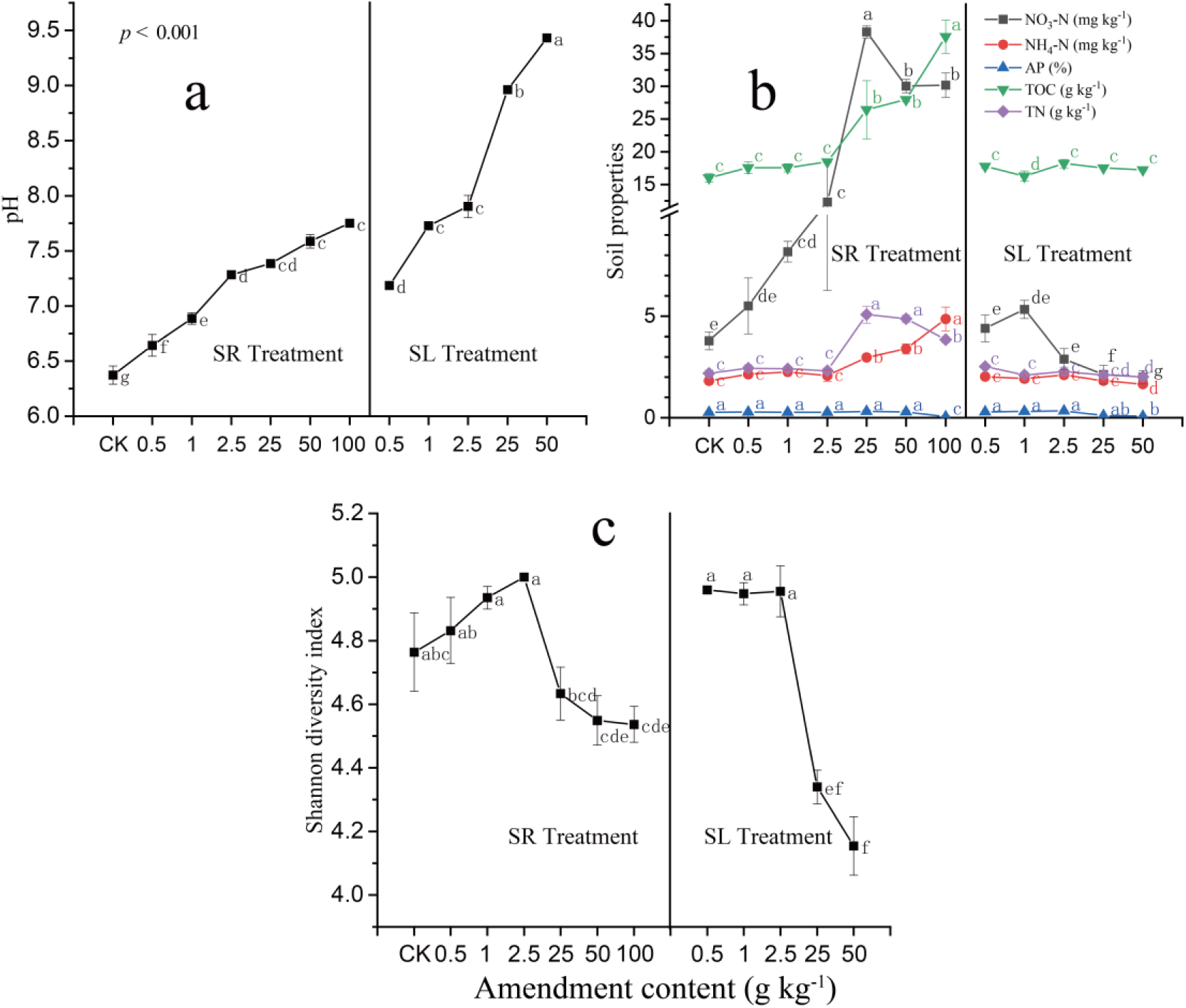
Variations of Soil pH (a), TOC and nutrients (b), and (c) Shannon diversity index of bacterial induced by the amendment of GAS residues (SR) and lime (SL) in acid soil after incubated for 120 days; where soil properties were indicated by color lines, TOC (Total Organic Carbon, green), NO_3_-N (nitrite, black), TN, (Total Nitrogen, purple) NH_4_-N (ammonia, red) and AP (Available Phosphorus, blue) (n=3).

Addition of GAS residues also increased soil carbon and soil nutrients—specifically, TOC, TN, NO_3_-N and NH_4_-N (Fig. 1b). TOC and NH_4_-N progressively increased as more GAS residues were added, increasing by 134.28% (TOC) and 167.80 % (NH_4_-N) with the amendment of 100 g kg^-1^ GAS residue. Soil nitrogen, TN and NO_3_-N showed peak values when 25 g kg^-1^ GAS residues were added, a threshold value prior to which proteins in the GAS residues decomposed or dispersed quickly, but after which anaerobic soils limited the activities of soil microbes and the transfer of proteins into small molecular and inorganic matter so that more NH4-N and NOx were produced and released into the air (27).

### 3.2 Amendment effects on bacterial diversity

Soil pH and nutrients are the important factors that can affect the soil bacterial community and diversity indicated by Shannon index. Amendment of GAS residues and lime both significantly affected soil bacterial diversity (Fig. 1c). Specifically, amendment of up to 2.5 g kg^-1^ GAS residues increased diversity, as measured by Shannon diversity index, from 4.76 (CK) to 4.99. The addition of more GAS residue decreased bacterial diversity; the addition 75 g kg^-1^ GAS residues resulted in the lowest Shannon diversity index value (4.55) across treatments. Similar to GAS residue, the addition of lime had a positive effect on diversity, but had a negative effect when more than 2.5 g kg^-1^ was added (Fig. 1c).

Previous studies have proposed that soil pH affects soil bacterial community structure and diversity (11, 28-30). N. Fierer and R. B. Jackson (10) found that the diversity and richness of soil bacterial communities differed by ecosystem type, and these differences could largely be explained by soil pH (*r*^2^ = 0.70 and *r*^2^ = 0.58, respectively; *P* < 0.0001 in both cases). Bacterial diversity was highest in neutral soils and lower in acidic soils, with soils from the Peruvian Amazon the most acidic and least diverse in their study (10). Our results were consistent with the observations reported in biochar amendment by Q. Li et al. (31), which found that bacterial diversity was relatively low in alkaline soils. We found that soil bacterial diversity increased with the additions of GAS residues and lime amendments between 0.5-2.5 g kg^-1^, likely because of the shift from acidic to neutral soil pH. Bacterial diversity declined when more GAS residue or lime were added (more than 25 g kg^-1^), likely due to different factors. For the SL (lime) treatment, high soil pH likely restricted the growth and reproduction of soil bacteria. This is consistent with the results reported by J. Xiong et al. (32), in which soil pH was negatively correlated with soil bacterial diversity in alkaline lake sediments across the Tibetan Plateau. Nevertheless, for the SR (GAS residues) treatment, high soil nutrients, such as NO_3_-N and NH_4_-N, likely contributed to the simplification of bacterial diversity. These results were contrast to the previous work, in that the addition of N had no significant effect on bacterial diversity while elevated P could increase bacterial diversity marginally (33).

However, pH was not likely responsible for the decline of soil bacterial diversity in the SR (GAS residue) treatment, because the peak value of soil pH in the SR treatment was equal or less than 7.8, the pH that yielded maximum bacterial diversity in the SL treatment. Instead elevated soil nutrients likely caused declines in bacterial diversity in the SR treatment. Soil nutrient concentrations were significantly negatively correlated with bacterial diversity index (Table S1). For example, TOC, TN, NH_4_-N and NO_3_-N were all negatively correlated with Shannon (*p* < 0.05) index. B. J. Campbell et al. (14) and A. Koyama et al. (34) reported similar negative relationships between bacterial diversity and N additions in Arctic tundra soils. Our results contrast, however, with other studies that show positive relationships between soil nutrients and bacterial diversity (33), which could be caused by differences in the types of soil, and the types and amount of nutrients amended (10, 35, 36).

### 3.3 Amendment effects on bacterial community composition

Our assessment generated 2016767 high-quality 16S rRNA gene sequences. The bacterial community composition of the soil samples shifted significantly as a result of the amendment of GAS residues and lime, and were clearly distinguished (Fig. 2a, ANalysis Of SIMilarity (ANOSIM), R = 0.823, *P* < 0.001) among the different amendment treatments, and such a pattern was confirmed by hierarchical clustering based on genus (Fig. 2b). Specifically, the analysis identified three groups of bacterial community compositions (*i.e*., in soils amended with high concentrations of lime, soils amended with high concentration of GAS residues and soils amended with low concentrations of lime or GAS residues). The reason contributed to these changes may attribute to elevated high soil pH and high nutrients availability induced by over amended lime and GAS residues.

**Fig. 2.**
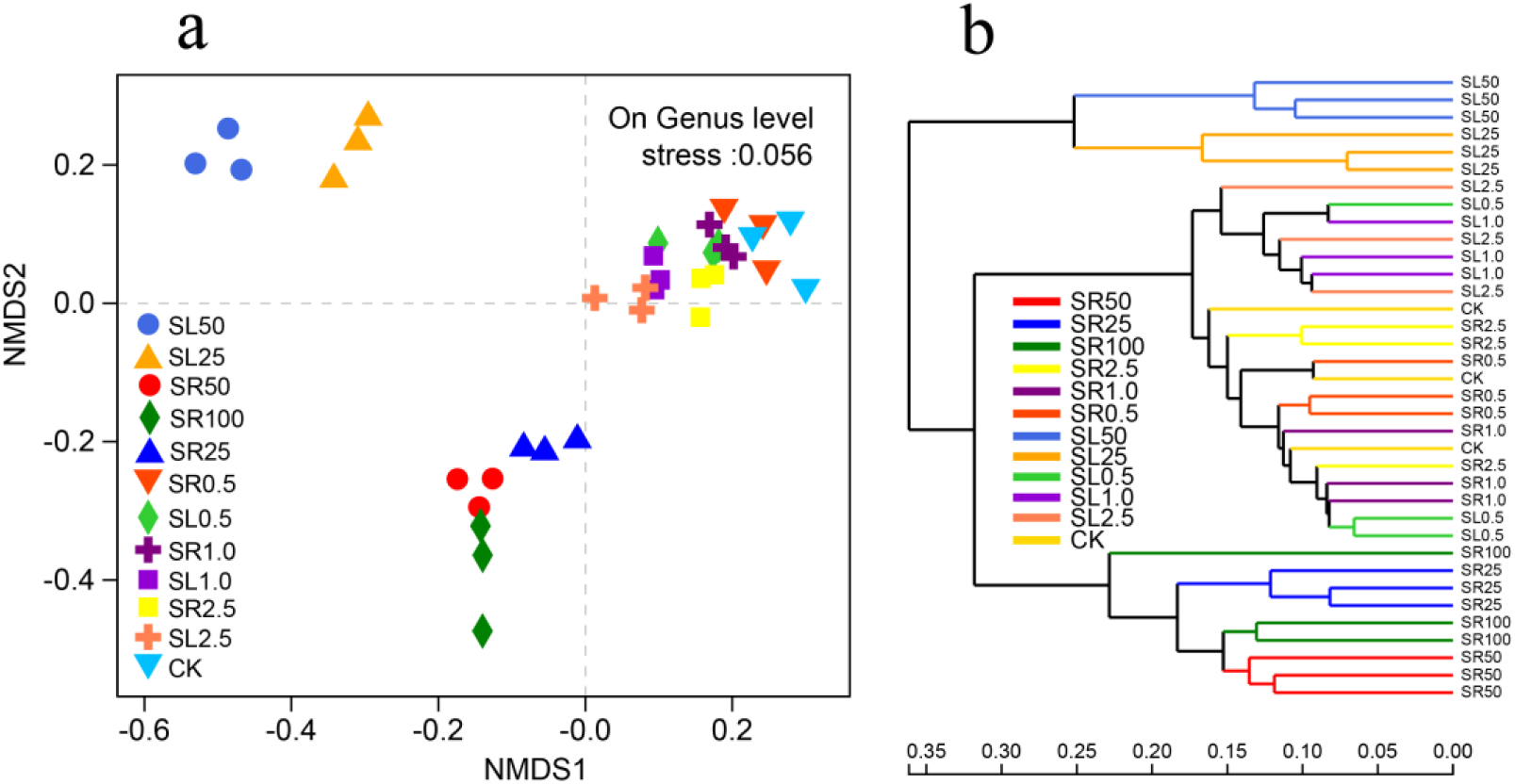
Non-metric multidimensional scaling (NMDS) of soil bacterial communities based on 16S rRNA within plots as affected amendments (a); and hierarchical clustering of treatments (b). SR represents GAS residues treatments, SL represents lime treatments and the numbers 0.5, 1, 2.5, 25, 50 and 100 represent the amendment levels (g kg^-1^), CK represents the control (n=3)

Amendment of GAS residues and lime induced different changes in bacterial community compositions. The most abundant bacterial phyla detected in soil samples were Proteobacteria (about 22%), Firmicutes (range from 15-35%), Chloroflexi (range from 12% - 22%), Actinobacteria (range from 8% - 20%), Acidobacteria, Gemmatimonadetes, Cyanobacteria and Bacterioidetes (Fig. 3). The amendment of GAS residues significantly increased the relative abundance of Firmicutes, Gemmatimonadetes, Bacterioidetes and Deinococcus-Thermus, but significantly decreased the relative abundance of Chloroflexi, Actinobacteria, Acidobacteria, Cyanobacteria and Planctomycetes. In SL (lime) treatments, we detected similar decreases in the abundance of Chloroflexi, Acidobacteria and Planctomycetes, but in contrast to SR (GAS residue) treatments, the abundance of Cyanobacteria increased significantly (Fig. 3). Gemmatimonadetes and Bacterioidetes increased in relative abundance in SR treatments but not in SL treatments, possibly because Gemmatimonadetes and Bacterioidetes were responsible for the degradation of soil carbon and the emission of CO_2_ (37, 38). Amendment of GAS residues (especially high levels) increased soil TOC, which likely contributed to increases in relative abundance of Gemmatimonadetes and Bacterioidetes. Our results contrast with previous studies that found that high soil pH (with biochar addition) resulted in decreases in Firmicutes abundance (39). That result may reflect the integrated effects of elevated soil pH, C and N (particularly N), because Firmicutes is involved in N cycling (40).

**Fig. 3.**
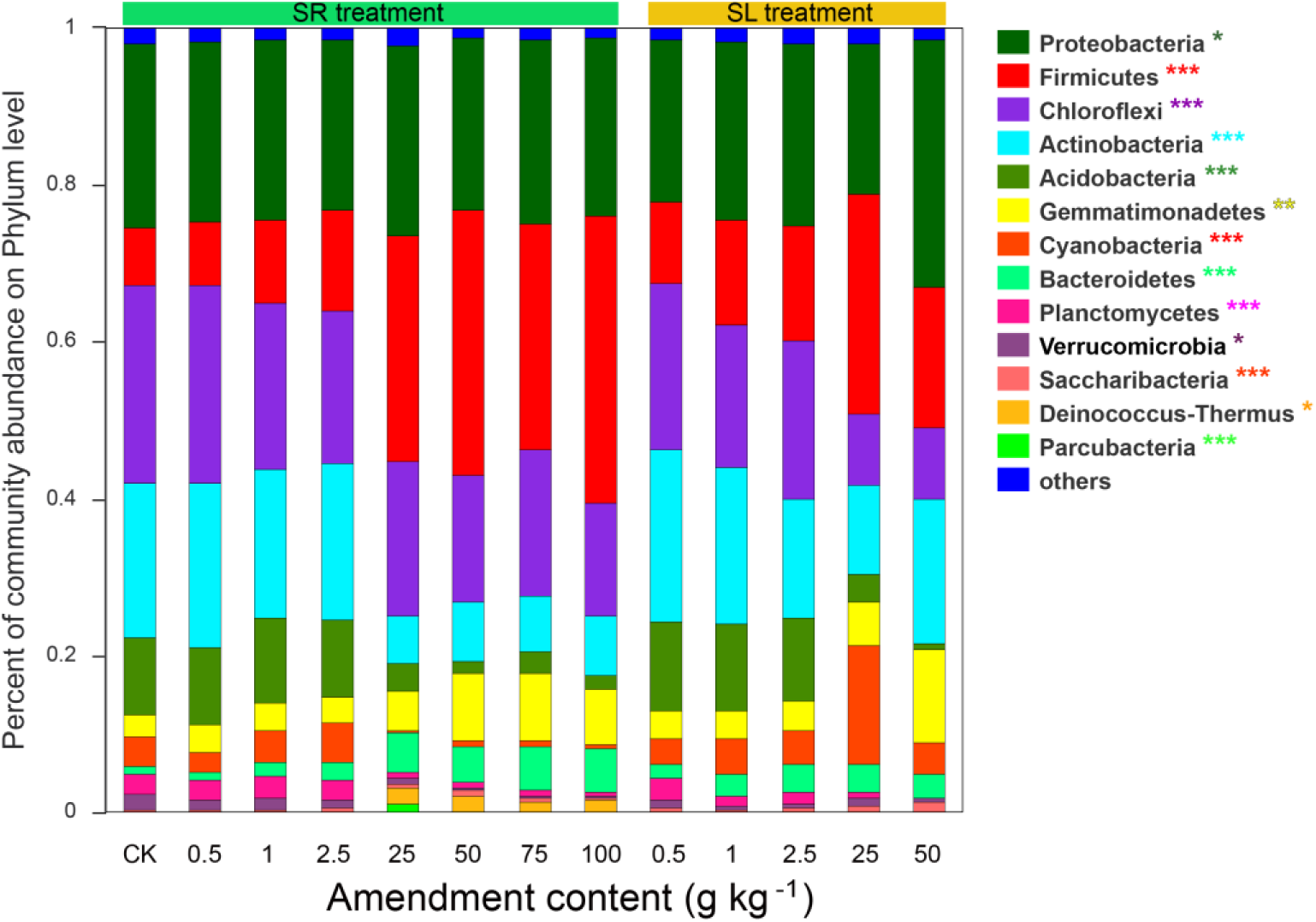
16S rRNA sequence-based microbial community composition of GAS residues (SR) and lime (SL) amended soils on the bacterial phylum level. Different colors indicate different phylum as showed in legend; and the symptoms *, * and *** represent significance at 0.05, 0.01 and 0.001 levels (n=3).

Significant changes of soil bacterial communities in SR (GAS residues) treatments can also be explained by copiotroph and oligotroph mechanisms driven by changes in soil nutrients. Copiotrophs preferentially consume labile soil organic C pools, have high nutritional requirements, and can exhibit high growth rates when resources are abundant. In contrast, oligotrophs exhibit slower growth rates and are likely to outcompete copiotrophs in conditions of low nutrient availability due to their higher substrate affinities (41). In our study, the amendments of GAS residues induced oligotrophic soil conditions that may have benefited copiotrophs and disadvantaged oligotrophs. Our results were similar to the previous studies, in that bacteria belonging to the Acidobacteria phylum were most abundant in soils with very low resource availability and their relative abundances declined in soils amended with high concentrations of organic C, N and P (33, 41).

### 3.4 Correlations between soil properties and bacterial community composition

Our results indicated that soil properties were significantly correlated with bacterial phyla (Fig. 4a) and genera (Fig. S1a) in GAS residues amended soils. For example, pH, NH_4_-N, NO_3_-N and TN had significant positive correlations with the relative abundance of Gemmatimonadetes, Tenericutes, Chlorobi, Firmicutes, Bacteroidetes and Deinococcus-Thermus (*p* < 0.01). In addition, significant negative correlations were detected with Antinobacteria, Cyanobacteria, Nitrospirae, Acidobacteria, Planctomycetes and Verrucomicrobia (*p* < 0.01). As previously discussed, Gemmatimonadetes, Tenericutes, Chlorobi, Firmicutes, Bacteroidetes and Deinococcus-Thermus are all involved in C and N cycling, and amendment of GAS residues (especially at high concentrations) induced high levels of C and N. The negative effects of SR treatment on Antinobacteria, Cyanobacteria, Nitrospirae, Acidobacteria, Planctomycetes and Verrucomicrobia may reflect sensitivities of some of these groups to higher soil pH; previous studies have found that some of these groups decrease in abundance after fertilizers are applied. For instance, D. R. Nemergut et al. (37) reported that the relative abundance of Verrucomicrobia declined in a fertilized soil. R. T. Jones et al. (42) reported that the abundance of Acidobacteria relative to other bacterial taxa was highly variable across soils, but correlated strongly and negatively with soil pH. B. J. Baker et al. (43) and N. Xu et al. (44) reported that Nitrospirae was active in nitrogen cycling, specifically as nitrite oxidation, and SR treatment may have increased labile nitrogen availability in the soil, possibly restricting growth of Nitrospirae. Although S. N. R. Prasanna (45) reported that Cyanobacteria prefer neutral to slightly alkaline pH for optimum growth, high nutrients may restrict its relative abundance.

**Fig. 4.**
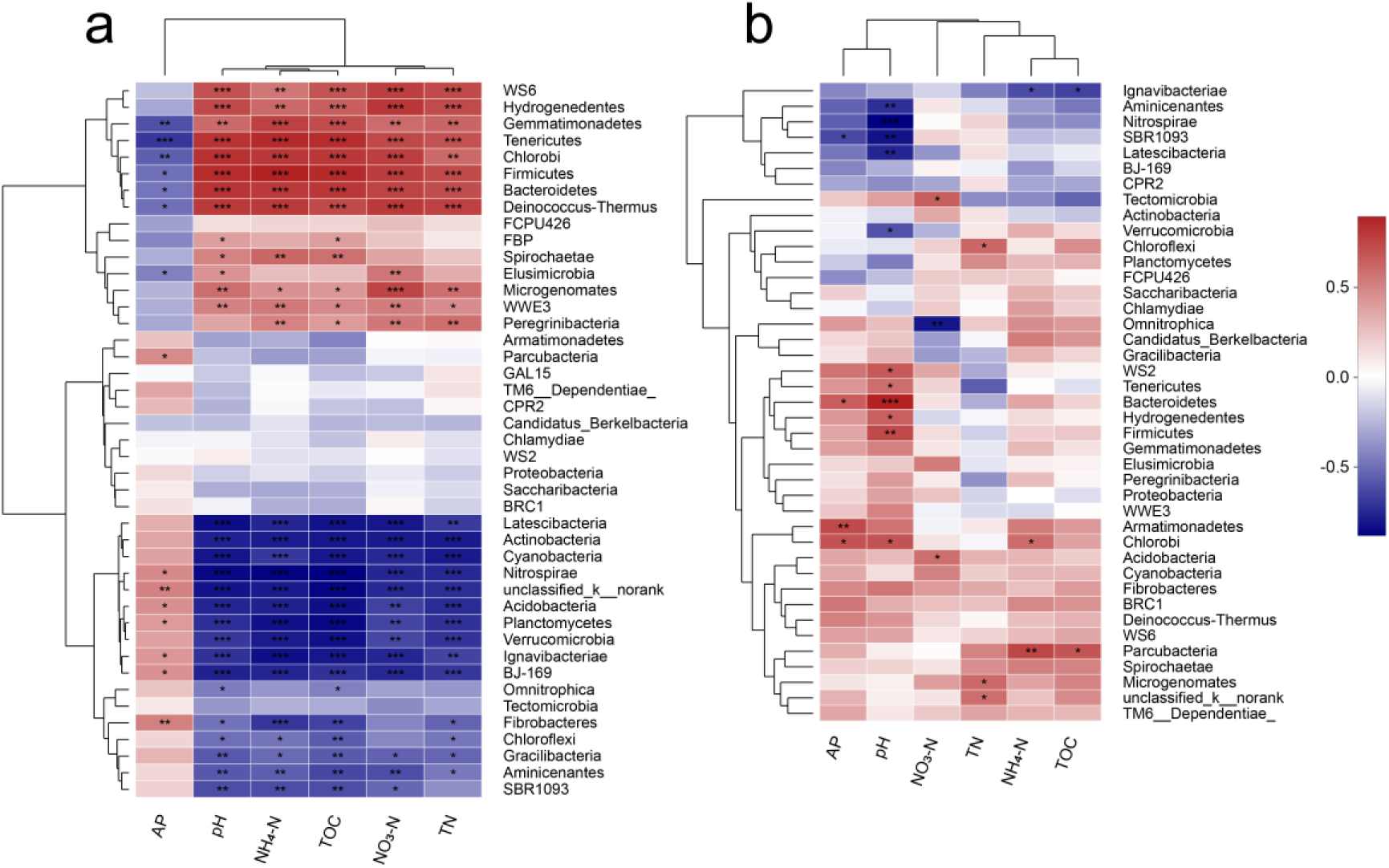
Spearman correlations between soil properties and bacterial community composition on phylum level after amendment of GAS residues (a) and lime (b); where blue and red colors represent negative and positive correlations,; TOC represents total organic carbon, TN represents total nitrogen, N-NH_4_^+^ represents ammonia, N-NO_3_^−^ represents nitrite and AP represents available phosphorus, and the symptoms *, ** and *** represent significant levels of 0.05, 0.01 and 0.001(n=3).

For the SR treatments, most significant correlations were found in soil pH; the only groups with relative abundance that was significantly positively related to pH were Bacteroidetes, Firmicutes and Chlorobi with soil pH (Fig. 4a). Fewer phyla showed significant correlations with soil nutrients in the SL treatments compared to the SR treatments. These results suggest that more phyla were affected by C, N (*ie*., TOC, NH_4_-N, NO_3_-N and TN) in SR treatments. In SR treatments, which increased both soil pH and nutrient contents, soil pH does not appear to be a good predictor of bacterial community composition and diversity.

To evaluate the main factors driving changes in microbial communities in our experimental treatments, we used Non-metric multidimensional scaling (NMDS) of soil bacteria community composition based on Bray-Curtis distances linear regression (Fig. 5). In SL treatments, soil pH was the factor most strongly related to changes in bacterial community composition (*R*^2^ = 0.90) (Fig. 5a). However, in SR treatments soil nutrients tend to shape changes in bacteria composition (Fig. 5b). Our results are consistent with previous research that included a sub-set of the soils included in this survey; they found that the abundances of Bacteroidetes, Betaproteobacteria, and Acidobacteria were most strongly related to estimated carbon availability, not soil pH (41). In addition, soil nutrients (N and P) additions could significantly affect soil bacterial community compositions were also confirmed (33). Nevertheless, our results contrast with other studies that suggest pH as a strong predictor of bacterial community composition and diversity (11, 28, 46).

**Fig. 5.**
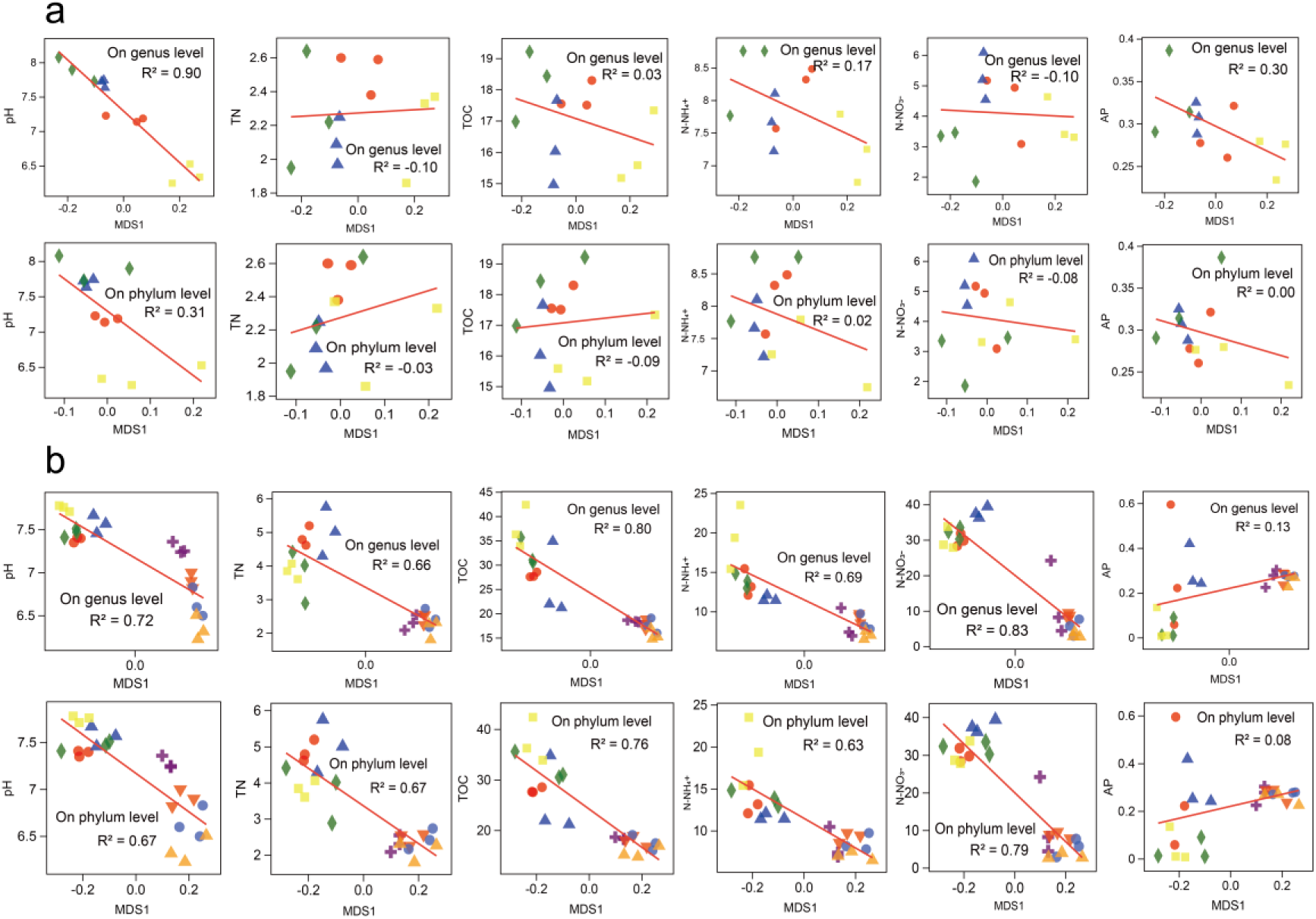
Soil properties and Non-metric multidimensional scaling (NMDS) of soil bacteria community composition based on Bray-Curtis distances linear regression after amending with lime (a) and GAS residues (b); TOC represents total organic carbon, TN represents total nitrogen, N-NH_4_^+^ represents ammonia, N-NO_3_^−^ represents nitrite and AP represents available phosphorus (n=3).

Apart from determining diversity another primary goal of comparing microbial communities is to identify specialized communities in samples. Groups were shown in cladograms, and LDA scores of 2 or greater were confirmed by LEfSe (Fig. S2). In GAS’s residues treated soils, 21 bacteria phyla were enriched dominated with Actinobacteria, Bacteroidetes and Firmicutes followed by Acidobacteria, Cyanobacteria and other 16 phyla. Firmicutes were most abundant in the GAS residues 50 g kg^-1^ treatment, which may have been caused by the high soil pH; it was the same pH as in the lime 2.5 g kg^-1^ treatment. To evaluate the effects of amendments on shaping microbial community composition, we compared results from high levels of the GAS residues amendments and low levels of lime amendments with results from low levels of GAS residues amendments, because high levels of GAS residues amendments and low levels of lime amendments showed similar effects on soil pH. The results showed that more specialized bacteria phyla were detected in soils treated with high levels of GAS residues amendments, which suggests that GAS residues can shape bacterial community compositions.

### 3.5 Correlations between enzyme activities and bacterial community composition

Four enzymes related to C, N and P cycling in soil were determined to uncover the status of soil nutrients and C cycling after the amendment of both GAS residues and lime. We conducted Spearman Correlation analysis to identify the relationships between microbial communities and soil enzyme activities, and significant correlations were found in SR treatment (Fig. 6). For example, we found that β–D– cellobiosidase (CB), β–1,4–N–acetylglucosaminidase (NAG) and β–1,4–glucosidase (BG) activities were significantly correlated with microbial communities both on phylum (e.g., Bacteroidetes, Deinococcus-Thermus, Gemmatimonadetes, Actinobacteria, Acidobacteriae and Nitrospirae) level (Fig. 6a) and genus level (Fig. S3a) in soils treated with GAS residues amendment. Many fewer correlations were found in soils treated with lime amendment (Figs. 6b, S3b). The strong correlations in soils treated with GAS residues may be caused by its influence on substrate such as C and N. M. Waldrop et al. (47) reported that bacterial community composition was correlated with BG, ACP, and sulphatase activity, and concluded that enzyme activity may provide a useful linkage between microbial community composition and carbon processing. As Fig. 6a shows, significant and negative correlations between bacterial phyla such as Actinobacteria, Acidobacteriae and Nitrospirae, and enzyme activities such as CB NAG and BG. Previous studies suggested that N additions resulted in significant reductions in soil microbial activity (13), diversity (14) and community structure composition (15) because of increases in C sequestration and/or decreases in soil respiration rates (13). However, in our study, enzyme activities significantly increased after the addition of GAS residues, which contain abundant N in the form of proteins and amino acids. It is possible that the elevated organic C and N induced by GAS residues treatments promoted hydrolyzation but suppressed the mineralization the C and N (48, 49). Our results are consistent with the results reported by M. Carreiro et al. (50), who found that microbes responded to N by increasing cellulase activity.

**Fig. 6.**
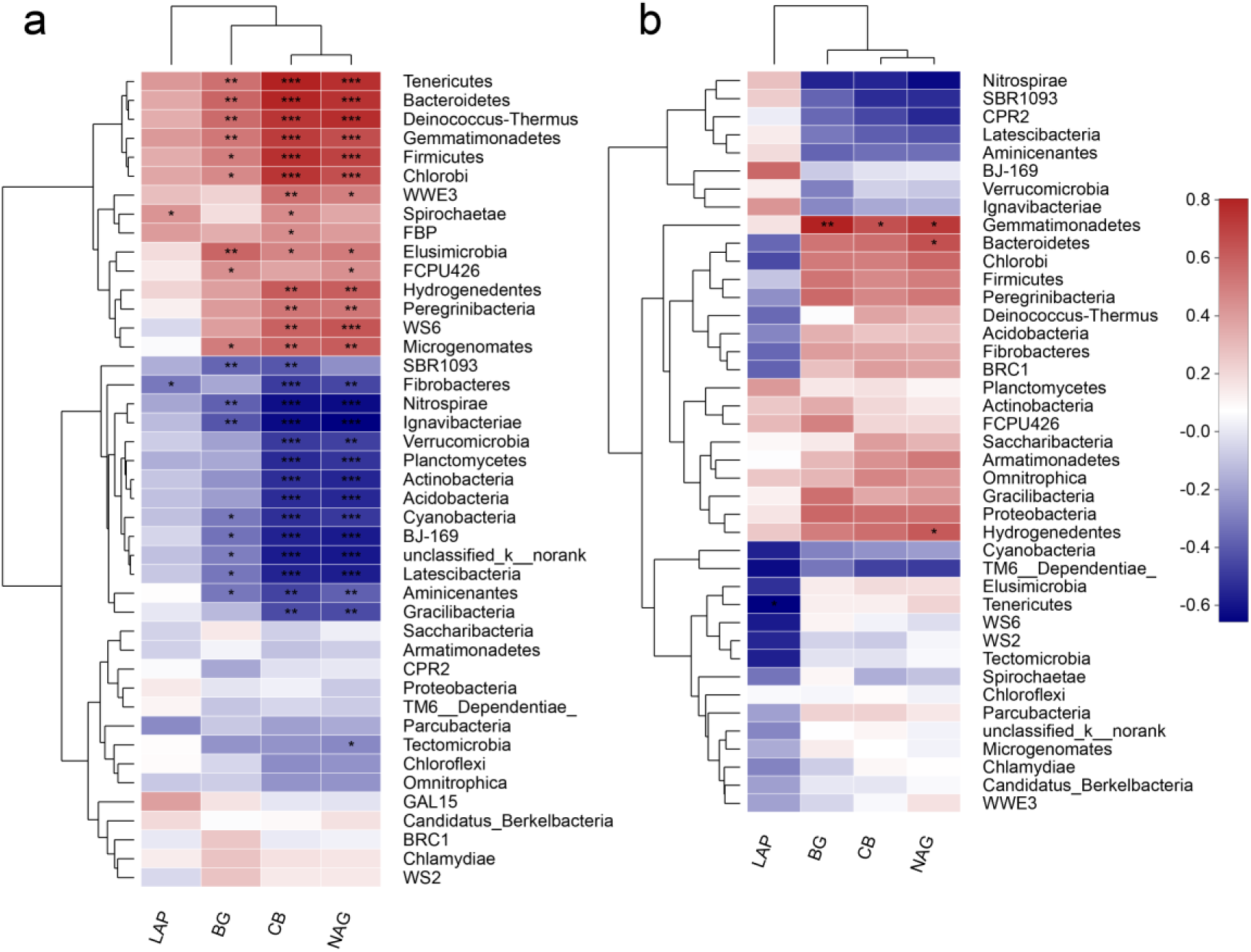
Spearman correlations between soil enzyme activities and bacterial community composition on phylum level after amendment of GAS residues (SR) (a) and lime (SL) (b); where blue and red colors represent negative and positive correlations; β–1,4– glucosidase (BG), acid phosphatase (ACP), β–1,4–N–acetylglucosaminidase (NAG) and β–D–cellobiosidase (CB); the symptoms *, ** and *** represent significant levels of 0.05, 0.01 and 0.001 (n=3).

Overall, in this study, the amendment of GAS residues significantly increased soil pH due to the CaCO_3_ component of the GAS’s shell. However, GAS residues had weaker effects on soil pH than did lime treatments. For example, at the same amended levels, the pH of soils amended with lime increased sharply, while soils amended with GAS residues rose only to near-neutral pH. In addition, amendment of GAS residues resulted in increased levels of soil nutrients, which could in turn lead to increased bacterial diversity at low amendment levels and decreased bacterial diversity at high amendment levels. That likely attribute to the amendment of GAS residues induced a copiotrophic environment in which, the relative abundance of copiotrophic bacterial communities were increased while oligotrophic bacterial communities were reduced. What’s more, soil pH also responsible for the changes of soil bacterial communities. For instance, Gemmatimonadetes, Tenericutes, Chlorobi, Firmicutes, Bacteroidetes and Deinococcus-Thermus were all increased with the addition of GAS residues. Some of them such as Gemmatimonadetes, Tenericutes, Chlorobi and Bacteroidetes were increased due to their roles in C and N cycling, while some of them were decreased because they were suppressed at a higher pH environment. Most researchers suggested that pH was the best predictor for bacterial diversity and community compositions across different types of land-use. Nevertheless, in our study, soil pH may not be the best predictor of bacterial community composition or diversity; rather soil nutrients (*ie*., NH_4_-N and NO_3_-N) and soil TOC showed stronger correlations with bacterial communities. That likely because the amendment of GAS residues induced elevated soil pH and nutrient content at the same time. Compared with lime treatment, the amendment of GAS residues caused more enriched bacterial phyla, that likely due to the soil nutrients difference between GAS residues treatment and lime treatment. Considering the nutrients cycling, soil enzymes activities related to C, N and P were determined and analyzed, our results suggested that soil enzymes activities showed similar correlations to bacterial communities with soil nutrients. Those indicated that the increases in enzymes activities were attributed to the elevation of soil nutrients induced by the amendment of GAS residues.

## 4 Conclusion

Our study proposed that GAS residues may be appropriate to remediate acidic soil, improve soil quality and reduce GAS populations in areas subject to GAS invasion. In practice, it may not be practical to dry and crush GAS into powder before applying it to soils. Instead, practitioners could create GAS residues at lower costs by collecting living or dead GAS, spreading them on the soil surface and smashing them using high speed rotary tiller. Also, we suggest applying GAS residues to nonirrigated farmland to reduce the potential water pollution. We suggest amending GAS at 2.5 – 25 g kg^-1^, which appears to be better for soil health and bacterial diversity. These recommendations warrant further testing in the field, but results of our greenhouse experiments suggest they hold promise.

## Acknowledgement

This work was supported by the National Natural Science Foundation of China (No. U1701236, 31870525, 41871034), Guangdong Modern Agricultural Technology Innovation Team Construction Project (No. 2018LM1100) and Guangdong Provincial Key Laboratory of Eco-Circular Agriculture (No. 2019B030301007). We would like to thank Dr. Abe Miller-Rushing for his assistance with English language and grammatical editing of the manuscript.

